# Genetic Variability and Correlation Coefficients of Major Traits in Early Maturing Rice under Rainfed Lowland Environments of Nepal

**DOI:** 10.1101/520338

**Authors:** Dev Nidhi Tiwari, Santosh Raj Tripathi, Mahendra Prasad Tripathi, Narayan Khatri, Bishwas Raj Bastola

## Abstract

Genetic variability is the fundamental requirement of any crop breeding program to develop superior cultivars. The objective of this study was to estimate the genetic variability and find out the correlation among the different quantitative traits of rainfed early lowland rice. The experiment was conducted consecutively two years during 2015 and 2016 in wet season across the four different locations in Regional Agricultural Research Station, Khajura, National Wheat Research Program, Bhairahawa, National Maize Research Program, Rampur and National Rice Research Program, Hardinath along the Terai region of Nepal representing sub-tropical agro-climate. Seven genotypes including Hardinath-1 as standard check variety were evaluated in the randomized complete block design with three replications. Various quantitative traits were measured to investigate the variability and correlation coefficients. All the genotypes and locations showed significant variations for all the traits considered. Genotypic coefficient of variation was lower than phenotypic coefficient of variation for all traits studied. The magnitudes of genotypic coefficient of variations were relatively higher for grain yield, 1000-grain weight and days to heading. The highest broad sense heritability of 94% was recorded in days to maturity and the lowest heritability of 16% was observed in plant height. Positive and highly significant correlations were found both in genotypic and phenotypic levels between days to heading and days to maturity (r_g_=0.9999**, r_p_=0.997**), days to heading and grain yield (r_g_ =0.9999**, r_p=_ 0.9276**), days to maturity and grain yield (r_g_ =0.9796**, r_p_=0.9174**). However, negative and highly significant genetic correlation was observed between plant height and 1000 grain weight (r_g_ = -0.9999**). Thus results indicated that days to heading, days to maturity, grain yield, 1000 grain weight demonstrated higher heritability and remarkable genetic advance could be considered for the most appropriate traits for improvement and selection of trait to achieve stable and high yielding early rice genotypes under rainfed environments.

## INTRODUCTION

Rice (*Oryza sativa* L., 2n=2x=24) is the predominant staple food crop in Nepal and ranks first in position in terms of production and productivity thus contributed significantly to livelihood of majority of people (Tiwari et al., 2018). It is grown in 1.36 million hectare and producing 4.3 million tons with productivity of 3.15 t ha^-1^ (MoAD, 2017). The several studies reported that about 2000 rice landraces exist in Nepal that are assumed to be grown from 60 to 3050 m altitude. It also reported that Nepal is one of the centers of rice diversity (Joshi, 2005). National Rice Improvement Program was established in 1972 at Parwanipur to organize the research and development works on rice as a major commodity crop that has significant contribution to the food security and economy of the nation (Joshi, 2005, MN Paudel, 2017). The systematic and deliberate efforts from Nepal Agricultural Research Council (NARC) have lead to release 81 rice varieties. These varieties aimed for different production ecosystems comprising terai, river basin, low hills, mid hills and high hills. Nepal’s production ecosystems have been broadly classified into two major ecosystems namely irrigated ecosystem, rainfed which comprises 49% and 51% respectively. Rainfed ecosystem constitutes integral component in Nepalese agriculture also has been further categorized as rainfed lowland and rainfed upland. Majority of agricultural lands in Nepal fall in the rainfed lowland ecosystem which accounts more than 55% and remaining areas are uplands.

In rainfed systems, drought is the major limiting abiotic stress that reduces productivity by 13–35% (Rosegrant et al., 2002). Yield is very low in rainfed rice compared to favorable growing conditions. Early or late maturing rice genotypes could be potential alternative to overcome the drought effects under rainfed rice growing conditions. The development of rice cultivars with improved drought resistance is thus an key element to alleviate risk, increased productivity and reduced yield variability in the rainfed environments (Suji et al., 2012). Around 90% of the world’s rice production is grown in the Asian continent and constitutes a staple food for majority of people worldwide (Khush, 2005). However, about 15% of Asian rice area experiences frequent yield loss due to drought which is widespread phenomenon of rainfed ecosystem (Kumar et al., 2007). Drought and upland areas occupy 31% and 11% respectively of the global rice-growing area which clearly indicated that drought is a major stress that limits rice production mainly in rainfed ecosystems (Kamoshita et al., 2008). To overcome yield reduction under rainfed and upland conditions, breeding objective should be targeted for development of drought tolerant cultivars.

The genetic improvement of any breeding population mainly depends upon the amount of genetic variability present. The genetic characters are mostly governed by poly genes which are highly influenced by the environment. Heritability of a genetic trait is important in determining the response to selection. In a similar manner high genetic advance coupled with high heritability offers the most effective condition for selection for a specific character (Islam et al., 2016). In order to understand the genetic variability of yield contributing attributes, relationship among them and their relation with yield are prerequisite to execute any breeding programme. The correlation coefficient may also help to know characters with little or no importance in the selection programme.(Singh et al., 2014).

The modern breeding program encompasses creating genetic variability, selection and utilization variation found in selected genotypes to generate new breeding materials. To reveal the genetic control of the any trait heritability, variance component and genotypic and environmental variance are key parameters of interest unraveling the gene action governing the desired trait. In order to initiate the efficient breeding program, information on estimation of variability and heritable elements of traits of particular genetic material is utmost necessary. Phenotypic and genotypic components of variation, heritability as well as correlation would provide valuable information for breeding of desirable traits (Bekele et al., 2013). Genetic variability among traits is essential for breeding and in selecting desirable genetic material (Atlin, 2003). Heritability, a measure of the phenotypic variance that is attributed due to genotype, has predictive function of breeding crops (Songsri et al., 2008). Generally, heritability represents the effectiveness of selection of genotypes that could be based on phenotypic performance (Bitew, 2016). Different genetic variability parameters, namely, Genotypic Coefficient of Variability (GCV), Phenotypic Coefficient of Variability (PCV), heritability and genetic advance for yield attributing traits is a major concern for any plant breeder for crop improvement programs. Similarly, information on correlation coefficients between grain yield and its component characters is essential since grain yield in rice is a complex character and is highly influenced by several component characters (Hossain et al., 2015). Likewise, correlation coefficient is another fundamental tool showing relationships among independent characteristics (Sravan et al., 2012).

The present study was carried out with an objective to understand the genetic background and inheritance of the different traits in rainfed early rice genotypes. Furthermore, the specific objective of this study is to estimate genetic variability, heritability and correlation among the various quantitative traits of rice grown under rainfed environments in order to aid the effective selection for successful breeding program. The findings of this study would help to identify the highly suitable genetic material and assist to design the subsequent breeding program to foster the varietal improvement programs.

## MATERIALS AND METHODS

### Experimental site and plant materials

Field experiments were carried out in wet season (Kharif) during 2015 and 2016 which is considered as main rice growing season in the South and Southeast Asia. Coordinated Varietal Trial (CVT) along the sub-tropical region which represented Terai region of Nepal. The collaborative research as multi-environment trial was conducted in four different research stations of the country. These four research stations included in the experimental study located from mid western to central Terai region of Nepal representing sub-tropical climatic domains.

### Description of Experimental Sites

#### Regional Agricultural Research Station (RARS), Khajura, Banke

RARS Khajura is located at Banke district under Province 5 and geographically it lies between 81°37’E longitude and 28°06’N latitude with an altitude of 181 meters above mean sea level. The average annual rainfall of the station is 1000-1500mm. However, delayed onset and early cessation of monsoon is characteristic feature of this station. The maximum and minimum temperature at the station is ±46°C and 5.4°C respectively, relative humidity ranging between 27-94%. The humidity remains low for most of the duration of the year. The soil of the station is sandy to silty loam with poor in organic carbon and available nitrogen but medium in phosphorus and potassium availability with pH varies from 7.2-7.5 (Source: Annual Report, RARS, Khajura, 2012)

#### National Wheat Research Program (NWRP), Bhairahawa, Rupendehi

NWRP Bhairahawa a is located at Rupendehi district under Province 5 and geographically it lies between 83°25’E longitude and 27°32’N latitude with an altitude of 104 meters above mean sea level. The maximum mean temperature at NWRP during the year was in month of May (37.2 ° C) and minimum mean temperature (10.5 ° C) in month of January. Total rainfall was 1879.7 mm with highest rainfall of 940.8 mm in the month of July (Source: Annual Report, NWRP, Bhairahawa, 2016). The relative humidity lowest (57%) in April and highest (85.4%) in September 2016.

#### National Maize Research Program (NMRP), Rampur, Chitwan

NMRP Rampur is situated at Chitwan district under Province 3 and geographically it lies between 84°19’E longitude and 27°40’N latitude with an altitude of 228 meters above mean sea level. It has humid climate with cool winter and hot summer. The soil is acidic with pH ranging from 4.6 to 5.7 and light textured and sandy loam soil. Average annual rainfall was 263.8 mm. Maximum and minimum temperature of 34.3°C and 6.14°C during month of August and January respectively. The relative humidity remains in between 86.8 to 97.8 % all the year round (Source: Annual Report, NMRP 2016).

#### National Rice Research Program (NRRP) Hardinath, Dhanusha

NRRP Hardinath is located at Dhanusha district in Province 2. It lies at the latitude of 26° 49’ E and at an altitude of 93 meters above from average mean sea level. The predominant soil type of the station is silty clay to sandy loam and pH is 6.3. The area has a subtropical climate with the average annual rainfall of 1281 mm. Maximum precipitation occurs in July and 80% of the total annual rainfall comes between June and September. The average relative humidity remains in between 55 to 90% around the year (Source: Annual Report, NRRP 2016).

The experimental materials consisted of promising early rice breeding lines received from International Rice Research Institute (IRRI, Philippines). These lines were evaluated in International Rainfed Lowland Observation Nursery (IRLON) under rainfed environment and superior lines were selected from this preliminary stage of evaluation. Those selected lines further screened in initial evaluation trial (IET) and the genotypes observed as superior and early were promoted to coordinated varietal trial (CVT). This is the standard and established method of evaluation of genotypes at different stages for varietal development process. Seven rice genotypes including a standard check variety Hardinath-1 were evaluated for two years across all four environments as multi-environment trial to identify the stable and adaptable genotypes across the locations and years.

### Experimental design and cultivation practices

The details of genotypes evaluated in the experiment have been depicted in the Table 1. The experiments were laid out in a randomized complete block design (RCBD) with three replications. The gross experimental unit was 5 m × 2 m (10 m^2^) and the net harvested area was 7.36 m^2^. Twenty five rows of 2m long with spacing of 20 cm x 20 cm row to row and plant to plant distance was maintained. Seedbed was prepared from finely pulverized soil then seeding was done by hand drilling method on first week of June in each year in all locations. Before seeding adequate FYM and fertilizers were applied at the time of land preparation following standard agronomic practices. Sufficient care was taken for plant protection measures, irrigation and weeding at the seedling stage to make seedlings healthy. Twenty five days old seedlings were transplanted in the main field. Experimental fields were well prepared as per the agronomic standards. Weeding was done two times based on the weed infestation. The crop was subjected to recommended package of agronomical and plant protection practices to obtain a healthy crop. Fertilizers were applied at the rate of 80: 40:30 N: P_2_O_5_: K_2_O kgha^-1^. Full dose of Phosphorus (P_2_O_5_) and Potassium (K_2_O) and half dose nitrogen (N) was applied at the time of field preparation. The remaining half dose of nitrogen was applied in two installments at tillering and before panicle initiation stage in 25 and 45 days after transplanting.

**Table 1.** Descriptions of rainfed lowland early rice genotypes tested at different locations during 2015 and 2016 (*NRRP, Hardinath, RARS, Khajura, NMRP, Rampur and NWRP, Bhairahawa) of Nepal

| Entry | Genotypes | Source | Origin |
| --- | --- | --- | --- |
| 1 | 08FAN2 | IRLON | IRRI |
| 2 | B11586-FMR-1-1-R-2-1-1 | IRLON | IRRI |
| 3 | Hardinath-1 | Released variety | Nepal |
| 4 | IR 10L 151 | IRLON | IRRI |
| 5 | IR 88965-39-16-4 | IRLON | IRRI |
| 6 | IR09L 270 | IRLON | IRRI |
| 7 | IR55423-01 | IRLON | IRRI |

\*NRRP= National Rice Research Program, RARS= Regional Agricultural Research Station, NMRP=National Maize Research Program and NWRP=National Wheat Research Program

### Data collection and analysis

Different quantitative characters such as days to heading and maturity were recorded on visual observations. Days to heading and maturity was recorded in days as well as plant height (cm) was taken by measuring the length of five individual plants from the soil surface to the tip of the panicle. In addition panicle length (cm), tillers per square meter (number) and 1000 grain weight in gram (g) was also taken into consideration. Grain yield per plot was recorded and converted into grain yield kg ha^-1^.

### Statistical Analysis

Analysis of variance (ANOVA) was carried out on the data to assess the genotypic effects and mean comparisons among treatment means were estimated by the least significant difference (LSD) test at 5% levels of significance. The analysis of variance performed using RCBD design to derive variance components derived using the software packages ADEL-R and META-R develop by CIMMYT, Mexico. Various genetic parameters such as genetic variance, phenotypic variance, genetic coefficient of variation (GCV), phenotypic coefficient of variation (PCV), heritability (H^2^), genetic advance (GA) and genetic advance as percentage of mean (GAM) were computed by using the error mean square for genotype, environment, G x E and error variance components. All the above variability measurements are depicted in the equation (1-10). Similarly, phenotypic and genetic correlation coefficient was computed with the software program META-R.

### Genotype ‒by – Environment Model Formulation

G x E Model

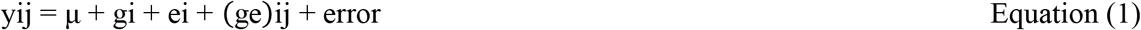

Where *yij* is the observed trait, *μ* is the grand mean, *gi* is the genotypic effect, *ei* is the environmental effect which is the two evaluation cycles and *(ge)ij* is the G x E interaction effect as proposed by (DeLacy et al., 1996).

### G x E phenotypic variance component model

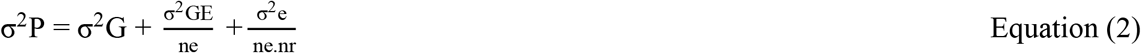

Where σ^2^P is the total phenotypic variance estimate, σ^2^G is the genotypic variance component, 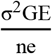 is G x E variance component and 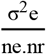 is the error variance component. The subscripts *ne* and *nr* implies number of environments and number of replications, respectively.

### Estimation of Genetic Parameters

The genetic parameters mainly genotypic variance (σ^2^g), phenotypic variance (σ^2^p), phenotypic coefficient of variation (PCV) and genotypic coefficient of variation (GCV) were derived by using the formula, suggested by (Burton and Devane, 1953) and (Johnson et al., 1955).

### Genotypic variance component

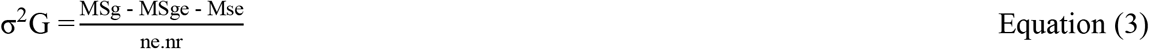

Where MSg represents mean square of genotypes, MSge is the genotype by environment mean square and MSe indicates error mean square.

### Genotype by Environment variance component

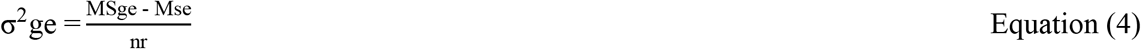

### Environmental variance component

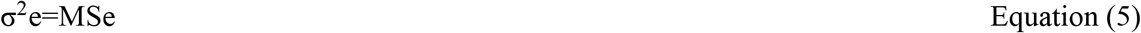

Genotypic coefficients of variation and phenotypic coefficients of variation were determined based on the method suggested by (Burton and Devane, 1953) as below:

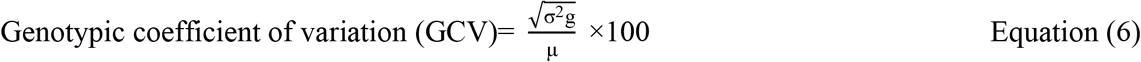

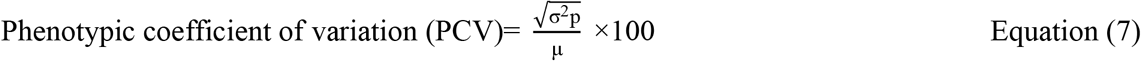

Where μ is the grand mean of the trait.

Broad sense heritability (H^2^) as percentage was derived for each character using variance components as explained by (DeLacy et al., 1996).

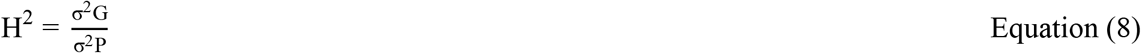

The heritability estimate was categorized as described by (Robinson et al., 1949) as below 0 - 30% = low, 30 - 60% = medium and >60% = high

### Estimation of Expected Genetic Advance from Selection

The genetic advance at selection intensity (k) at 5% (2.06) was derived by using following formula (Johnson et al., 1955):

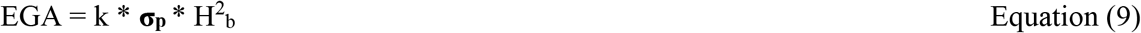

Where, EGA represents the expected genetic advance under selection; **σp** is the phenotypic standard deviation**; H^2^_b_** is heritability in broad sense and k is selection intensity

The genetic advance as percent of population mean was also derived by using the procedure of (Johnson et al., 1955).

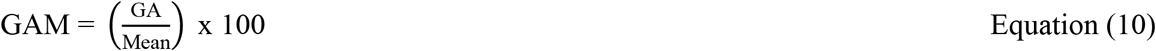

### Estimation of correlation coefficients

The Genotypic and phenotypic correlation coefficients between yield and yield attributing traits were computed as described by (Hossain et al., 2015) are depicted in equation 11 and 12 respectively.

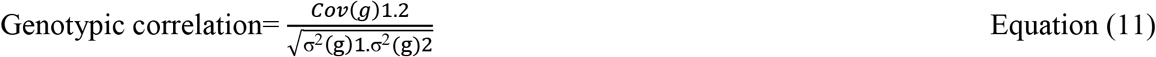

Where,

Cov(g) (xy)=Genotypic covariance between the variables X and Y
σ^2^ (g) 1=Genotypic variance of the variable X1
σ^2^ (g) 2=Genotypic variance of the variable X2

Similarly,

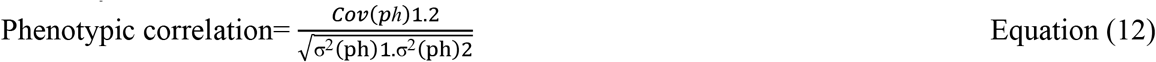

Where,

Cov (ph) (xy)=Phenotypic covariance between the variables X and Y
σ^2^ (ph) 1=Phenotypic variance of the variable X1
σ^2^ (ph) 2=Phenotypic variance of the variable X2

## RESULTS AND DISCUSSION

The analysis of variance exhibited the presence of significant differences among the tested genotypes for all characters indicating the existence of variability. The analysis of variance showed that the genotypes differed significantly (p<0.05) for plant height and highly significant (p<0.01) for days to flowering, days to maturity, thousand grain weight, grain yield and tillers m^-2^. Furthermore, locations effects were pronounced significantly for all the measured traits as well as location by year effect were also differed for all the traits except for days to maturity (Table 2).

**Table 2.** Mean genotypic performance of seven genotypes across all location over two years (2015 and 2016).

| Genotype | DH<br>(DAS) | DM<br>(DAS) | PH<br>(cm) | Till m<sup>-2</sup><br>(no.) | GY<br>(kg ha<sup>-1</sup>) | 1000-gwt.(g) |
| --- | --- | --- | --- | --- | --- | --- |
| 08FAN2 | 85 | 113 | 105.43 | 229 | 3388.09 | 22.07 |
| B11586-FMR-1-1-R-2-1-1 | 85 | 113 | 105.76 | 243 | 3557.92 | 19.96 |
| Hardinath-1 | 81 | 108 | 105.44 | 223 | 3078.77 | 21.41 |
| IR 10L 151 | 88 | 115 | 105.81 | 243 | 4227.04 | 21.12 |
| IR 88965-39-16-4 | 89 | 116 | 105.57 | 247 | 4198.99 | 22.64 |
| IR09L 270 | 89 | 117 | 106.62 | 240 | 3954.77 | 19.89 |
| IR55423-01 | 92 | 120 | 106.12 | 230 | 4266.67 | 21.63 |
| Grand Mean | 87 | 115 | 108.10 | 238 | 3810.3 | 21.24 |
| F-Test (Geno.) | ** | ** | * | ** | ** | ** |
| F-Test (Loc.) | ** | ** | ** | ** | ** | ** |
| F-Test (Geno. x Loc.) | ** | Ns | * | * | ** | ** |
| LSD<sub>0.05</sub> | 3.00 | 3.30 | 6.84 | 57.68 | 490.4 | 1.75 |
| MSE<sub>Error</sub> | 27.7 | 33.5 | 143.82 | 10237.87 | 740014.9 | 9.48 |
| CV% | 6.05 | 5.03 | 11.10 | 42.60 | 22.57 | 14.49 |

\* and \*\* indicates significance at 5% and 1% level of P-value. DH stands for days to heading, DM for days to maturity, DAS implies days after seeding, PH refers to plant height, Till m<sup>-2</sup> refers to tillers per square meter, GY refers to grain yield and TGW refers to thousand grain weight

The statistical analysis using ANOVA with RCB design indicated that Mean Squares Error (MSE) estimates (Table 3) showed highly significant differences (p≤0.01) among genotypes for days to heading, days to maturity, grain yield and 1000-grain weight. Similarly, significant difference (p<0.05) was observed among genotypes for plant height and tillers m^-2^. All the measured traits demonstrated significant environmental difference (p≤0.01) including both year and locations effect collectively referred to as environmental effects (Table 3). The highly significant G x E interactions (p≤0.01) was found for days to heading, thousand grain weights and grain yield (kg ha^-2^). However, non-significant interaction was noticed for days to maturity while the significant G x E interaction (p<0.05) for plant height and tillers m^-2^ was influenced by both environment and genotype. It is obvious from the above findings that there is existence of variability in the tested genotypes in great extent (Table 2 and Table 3).

**Table 3.**
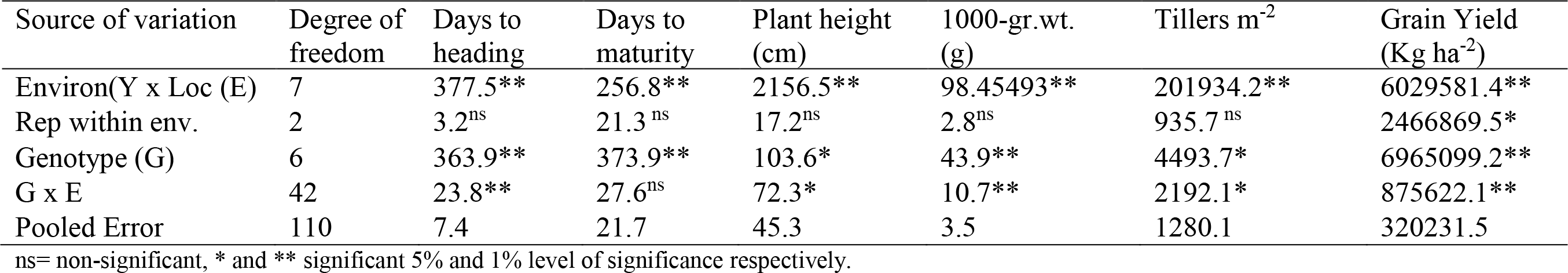
Genotype (G), environment (E) and G X E means squares for yield and yield components in rainfed lowland early rice genotypes

| Source of variation | Degree of freedom | Days to heading | Days to maturity | Plant height (cm) | 1000-gr.wt. (g) | Tillers m<sup>-2</sup> | Grain Yield (Kg ha<sup>-2</sup>) |
| --- | --- | --- | --- | --- | --- | --- | --- |
| Environ(Y x Loc (E) | 7 | 377.5** | 256.8** | 2156.5** | 98.45493** | 201934.2** | 6029581.4** |
| Rep within env. | 2 | 3.2<sup>ns</sup> | 21.3<sup>ns</sup> | 17.2<sup>ns</sup> | 2.8<sup>ns</sup> | 935.7<sup>ns</sup> | 2466869.5* |
| Genotype (G) | 6 | 363.9** | 373.9** | 103.6* | 43.9** | 4493.7* | 6965099.2** |
| G x E | 42 | 23.8** | 27.6<sup>ns</sup> | 72.3* | 10.7** | 2192.1* | 875622.1** |
| Pooled Error | 110 | 7.4 | 21.7 | 45.3 | 3.5 | 1280.1 | 320231.5 |

ns= non-significant, \* and \*\* significant 5% and 1% level of significance respectively.

The degree of variability for given trait is required for the improvement for breeding of field crops. The estimates of genotypic variation (σ^2^g), phenotypic variation (σ^2^p), genotypic coefficient of variation (GCV), phenotypic coefficient of variation (PCV), heritability (h^2^_b_) and genetic advance (GA), Genetic advance as percentage of mean (GAM) for different characters have been presented in Table 4. Phenotypic component of variance for all the measured traits was further partitioned into genotypic variance (σ^2^G), G x E variance (σ^2^GE) and error variance (σ^2^e). However, only genotypic variance was compared with total phenotypic variance to understand the magnitude of genotypic contribution to rice improvement.

**Table 4.** Estimation of means, ranges, variance components, phenotypic coefficient of variations, genotypic coefficient of variations, broad sense heritability, genetic advance and genetic advance as percent of mean of yield and yield traits of early maturing rice under rainfed lowland environment of Nepal.

| Traits | Mean | Range | s<sup>2</sup>g | s<sup>2</sup>p | PCV% | GCV% | H<sup>2</sup><sub>b</sub> | EGA | GAM |
| --- | --- | --- | --- | --- | --- | --- | --- | --- | --- |
| Days to heading | 86.9 | 81.1-92.2 | 13.86 | 14.81 | 4.42 | 4.28 | 0.94 | 7.40 | 8.50 |
| Days to maturity | 115.1 | 107.7-119.6 | 13.50 | 14.65 | 3.33 | 3.20 | 0.92 | 7.30 | 6.30 |
| Plant height (cm) | 108.1 | 105.4-106.6 | 0.58 | 3.59 | 1.75 | 0.70 | 0.16 | 0.62 | 0.58 |
| 1000-gr.wt.(g) | 21.2 | 19.9-22.6 | 1.24 | 1.68 | 6.10 | 5.24 | 0.74 | 1.97 | 9.30 |
| No. of Tiller m<sup>-2</sup> | 237.5 | 223.3-247.1 | 42.56 | 133.90 | 4.87 | 2.75 | 0.32 | 7.57 | 3.20 |
| Grain yield (Kg ha<sup>-2</sup>) | 3810.3 | 3078.7-4266.7 | 240385.20 | 276869.50 | 13.80 | 12.86 | 0.86 | 932.18 | 24.48 |

s<sup>2</sup>g = genotypic variance, s<sup>2</sup>p = phenotypic variance, PCV = phenotypic coefficient of variation, GCV = genotypic coefficient of variation, EGA=Expected genetic advance, GAM=genetic advance as percentage of mean.

The highest GCV of 12.86% was observed for grain yield followed by 5.24% and 4.28% GCV found in 1000 grain weight and days to heading respectively. However lowest GCV of 0.7% was recorded from plant height. On the other hand, highest phenotypic coefficient of variation (PCV) was observed in grain yield (13.8%) followed by 6.1% and 4.87% for 1000 grain weight and number of tillers m^-2^ respectively. The result showed that lowest PCV of 1.75% was observed in plant height similar to result found in case of GCV. This result is in consistent with finding reported by (Aditya and Bhartiya, 2013) in upland rice indicating selection for this trait is highly desirable. It was observed that phenotypic coefficient of variation was slightly higher than the genotypic coefficient of variation (GCV) for all the traits (Table 4). However, the difference between PCV and GCV was found very low reflecting very few influence of environment in the expression of characters or lower sensitivity of genotypes to environment and greater role of genetic control governing the character is in agreement with the results explained by (Sravan et al., 2012, Karim et al., 2007). The results obtained in our study is in agreement with the findings from previous works of (Bitew, 2016, Pandey and Anurag, 2010, Seyoum et al., 2012, Ubarhande et al., 2009, Hossain et al., 2015) suggesting environmental influence not noticeable in the expression of phenotypic characters. It means characters less affected by environment (Karad and Pol, 2008) and selection on the basis of phenotype independent of genotype could be effective for the improvement of such traits. Furthermore, studies also revealed that genotypes with high GCV of yield attributing traits are essential in breeding high yielding varieties through hybridization and selection. It is assumed that high degree of GCV give rise to variable offspring in the segregating generations (Bose et al., 2007).

The heritability (H^2^) estimate varied from 16-94% where highest heritability of 94% was recorded from days to heading and lowest 16% heritability observed in plant height (cm). Similarly, heritability of other traits was observed 92% for days to maturity, 86% for grain yield followed by 74% in case of 1000 grain weight. The H^2^ estimate for tillers m^-2^ was found low (34%) (Table 4). Similar findings also reported by (Seyoum et al., 2012) which mentioned moderately high heritability for days to maturity. Low heritability caused by high environmental influence over the expression of character.

The genetic advance as percentage of mean (GAM) showed that the highest GAM of 24.48 was observed in grain yield followed by 9.3 in thousand grain weights, 8.5 in days to heading, 6.3 in days to maturity and 3.2 observed in tillers m^-2^. GAM was found very low in plant height (0.58) (Table 4). In the present study, days to heading, thousand grain weight and grain yield showed higher heritability combined with high genetic advance as percent of mean (GAM) (Table 4). The finding is very closely supported by other workers suggesting additive gene action in the expression of character and hence selection would be successful (Pandey et al., 2009, Hossain et al., 2015, Bekele et al., 2013, Shukla et al., 2005, Suman et al., 2005).

Grain yield and thousand-grain weight showed moderately high heritability 86% and 74% coupled with high genetic advance in percent of mean 24.48 and 9.3 respectively. In tillers m^-2^ low to moderate heritability (32%) coupled with low to moderate genetic advance as percent of mean (3.2%) was observed. The present finding is in consistent with results from (Ullah et al., 2011, Seyoum et al., 2012) exhibiting the occurrence of additive gene effects and selection based on such traits would be highly desirable (Bose et al., 2007). Similarly, low heritability (16%) and its simultaneous low genetic advance as percent of mean (0.58%) were recorded in plant height reflected existence of non-additive gene action and greater environmental influence on the trait as investigated by (Akinwale et al., 2011, Bose et al., 2007). In this study, high heritability and genetic advance was observed in grain yield, days to heading and maturity indicated presence of additive genetic control also in agreement with (Islam et al., 2016, Bekele et al., 2013) during their study in aromatic and fine rice germplasm. In this study high heritability (92%) coupled with moderate genetic advance as percent of mean (6.3%) was recorded in the days to maturity. Such phenomenon suggested that these traits mostly governed by interaction of genetic and environmental components. Selection based on the genotype is not practical. Similar finding was reported by (Bekele et al., 2013) in days to maturity which is fully in agreement with the present finding. Moderate heritability (74%) with high genetic advance as percent of mean (9.3%) but low genetic advance of 1.97% was demonstrated by 1000-grain weight which is very interesting finding in this study as depicted in Table 4. This is highly consistent with the result obtained by (Karim et al., 2007, Kumar et al., 1998) which illustrated that selection for such trait could not bring desirable changes over the population mean.

Correlation coefficient analysis is widely used to measure the degree and direction of relationships between various traits including grain yield. Days to maturity possessed positive and highly significant correlation with days to heading at genotypic (r_g_ = 0.9999**) and phenotypic (r_p_ = 0.997**) levels. On the other hand, plant height exhibited positive and significant correlation with days to heading (r_g_=0.9359**) and maturity (r_g_=0.9999**) at genotypic level. However, phenotypic correlation was observed non-significant and positive (r_p_ = 0.5997, r_g_=0.6348) respectively with days to heading and maturity (Table 5). Days to heading and days to maturity showed significant and positive correlation at genotypic ((r_g_=0.9999** and 0.9796**) and phenotypic (r_p_=0.9276** and 0.9174**) levels respectively with grain yield as presented in Table 5. However, plant height had highly significant and negative genetic correlation (r_g_=-0.9999**) with 1000-grain weight. On the contrary it revealed non-significant and negative phenotypic correlation (r_p_=-0.4926) with thousand grain weight as given in Table 5. All the correlation of tillers m^-2^ with days to heading (r_g =_0.6462, r_p_=0.3623), maturity (r_g =_0.6642, r_p_=0.3877) and plant height (r_g =_0.512, r_p_=0.0539) was found positive but non-significant at genotypic and phenotypic levels presented in Table 5. Similarly, thousand grain weight had negative but significant genetic correlation (r_g =_-0.9999**). Besides this, correlation of this trait with days to heading (r_g =_0.0475), maturity (r_g =_0.0255) and tillers (r_g =_0.3028) at genotypic level was found non-significant and negative. There was non-significant but positive correlation between grain yield and thousand grain weight (r_g_=0.0725).

**Table 5.** Genotypic and phenotypic correlation coefficients of yield and yield attributing traits of rainfed lowland early rice genotypes

| **Traits** | **Type of correlation** | **DM** | **PH** | **Till m<sup>2</sup>** | **GY** | **TGW** |
| --- | --- | --- | --- | --- | --- | --- |
| **DH** | G | 0.9999** | 0.9359** | 0.6462 | 0.9999** | -0.0475 |
|  | P | 0.9969** | 0.5997 | 0.3623 | 0.9276** | 0.1479 |
| **DM** | G | 1 | 0.9999** | 0.6642 | 0.9796** | -0.0255 |
|  | P | 1 | 0.6348 | 0.3877 | 0.9174** | 0.1052 |
| **PH** | G |  | 1 | 0.592 | 0.7448 | -0.9999** |
|  | P |  | 1 | 0.0539 | 0.4141 | -0.4926 |
| **Till_m2** | G |  |  | 1 | 0.8404* | -0.3028 |
|  | P |  |  | 1 | 0.5710 | -0.0810 |
| **GY** | G |  |  |  | 1 | 0.0725 |
|  | P |  |  |  | 1 | 0.2955 |

“\*, \*\* Indicate significance at 0.05 and 0.01 probability levels”.
Where: GY = grain yield, DH = number of days to heading, DM = number of days to mature, PH = plant height, TGW = thousand grain weight and Till\_m2=tillers per square meter, G=Genotypic and P=phenotypic

The positive and significant correlation between grain yield and thousand grain weight was in consistent with the previous findings by (Islam et al., 2016). Thus significant and positive correlation of among the traits reflected that selection for one trait would directly affect the expression of other trait facilitating the selection and progress on breeding program. We found very similar findings in this study very consistent with the results from previous works in rice (Hossain et al., 2015). The highly significant but negatively correlation observed between plant height and 1000-grain weight indicated that selection for such negatively correlated traits would be very crucial affecting the progress under selection. Improvement in thousand grain weight character will lead to reduction of the grain yield or alteration in the days to flowering or maturity duration as well as change in plant height. The above statement is fully supported by report from (Newell and Eberhart, 1961) describing the situation when two characters show negative genotypic correlation it would be hard to apply simultaneous selection for these characters for development of a new variety. Therefore, judicious selection programme might be designed for simultaneous improvement of such component characters. The results revealed that the extent of genotypic correlation coefficients were higher than their phenotypic correlation coefficients indicating absence of environmental effects that enhanced genetic inherent associations (Sravan et al., 2012, Hossain et al., 2015). However, higher or equivalent phenotypic correlation coefficients corresponding to genotypic correlation coefficients suggesting that both environmental and genotypic correlation act in the same direction. Our finding is in agreement with the previous result in Aman rice presented by (Hossain et al., 2015).

## CONCLUSION

Phenotypic coefficient of variation (PCV) was found slightly higher than the genotypic coefficient of variation (GCV), higher heritability combined with high genetic advance as percentage of mean (GAM) and genotypic correlation coefficients were higher than their phenotypic correlation coefficients reflected that there is presence of additive genetic control and revealed absence of environmental effects that enhanced genetic inherent associations. The genotypes studied in this study possessed most valuable traits and thus should be incorporated in the breeding of lowland early rice varieties in the rainfed environments of Nepal.

## ACKNOWLEDGEMENTS

Author would like to provide sincere gratitude to NARC for providing financial source to undertake this study. I would like to express sincere thanks to all the collaborators from all stations as well as staffs of NRRP, Hardinath. We are very grateful to IRRI, Philippines for providing the new breeding lines which are suitable for rainfed environments. Finally efforts of all people who directly or indirectly supported to carry out this piece of work would be highly appreciated.

## DATA AVAILABILITY

All the relevant data files will be provided supporting the results obtained for this manuscript. I will be ready to submit them when necessary.

